# Bayesian neural networks for the optimisation of biological clocks in humans

**DOI:** 10.1101/2020.04.21.052605

**Authors:** G Alfonso, Juan R Gonzalez

## Abstract

DNA methylation is related to aging. Some researchers, such as Horvath or Levine have managed to create epigenetic and biological clocks that predict chronological age using methylation data. These authors used Elastic Net methodology to build age predictors that had a high prediction accuracy. In this article, we propose to improve their performance by incorporating an additional step using neural networks trained with Bayesian learning. We shown that this approach outperforms the results obtained when using Horvath’s method, neural networks directly, or when using other training algorithms, such as Levenberg-Marquardt’s algorithm. The R-squared value obtained when using our proposed approach in empirical (out-of sample) data was 0.934, compared to 0.914 when using a different training algorithm (Levenberg Marquard), or 0.910 when applying the neural network directly (e.g. without first reducing the dimensionality of the data). The results were also tested in independent datasets that were not used during the training phase. Our method obtained better R-squared values and RMSE than Horvath’s and Levine’s method in 8 independent datasets. We demonstrate that building an age predictor using a Bayesian based algorithm provides accurate age predictions. This method is implemented in an R function, which is available through a package created for predicting purposes and is applicable to methylation data. This will help to elucidate the role of DNA methylation age in complex diseases or traits related to aging.

## 1 Introduction

In recent years there has been an increased interest in studying the impact of methylation levels on aging. Some researchers, such as Horvath [1] and Levine [2], have developed biological clocks that are able to estimate the age of a patient by analyzing the level of methylation of blood cells and also tissues such as brain, breast or colon matter. At a chemical level, methylation is defined by the addition of a group methyl to cytosine base at 5-CPG-3 location, see Caifa and Zampieri [3], Patterson et al. [4], McBryan and Adams [5], Cerchietti and Melnick [6], Yuval and Cedar [7]. CPG represents the link between a Cytosine and a Guanine base by a phospodiester bond. It has been theorised by authors such as Lim and Maher [8] that methylation has an important role in regulating gene expression. However, the role of DNA methylation in the process of aging in humans remains unclear. Authors such as Jones [9] have noticed that “certain CPGs sites are highly associated with age, to the extent that prediction models using a small number of these sites can accurately predict the chronological age of donors”, referring to biological clocks such as those created by Horvath or Hannum. The impact of methylation on mortality has also been analyzed [10]. Horvath [11] found that methylation is impacted by some environmental and lifestyle factors, such as obesity. In particular, they found that obesity increases aging in the liver. Aging is clearly a complex process with many intertwined factors. For instance, the impact of telomere shortening in aging has been the object of several studies [12, 13, 14]. There have also been extensive studies regarding the link between methylation and cancer. For instance, Hashimoto [15] used methylation as a biomarker for gastrointestinal tumors and Pouliot [16] used a similar approach for breast cancer.

Although the reported prediction accuracy in both Horvath and Hannum’s age predictors was high, it can be improved by using better statistical methods. Both methods used Elastic Net (EN) technique, which aims to reduce the dimensionality of data. EN is based on a regularised regression that linearly combines the L1 and L2 penalties of the lasso and ridge methods. We hypothesise that using Bayesian neural networks (BNN) after dimensionality reduction seems a logical step because there is no indication that the level of methylation and the chronological age of the patient should follow a linear relationship. In addition, reducing the dimensionality of the data before applying neural networks is beneficial because it likely decreases the possibility of issues with local minimums, which is a frequently mentioned drawback of neural networks. It will also decrease the computational costs of the calculations required to train it. Therefore, our proposed method combines EN and BNN to predict chronological age based on methylation data. In particular, the dimensionality of the data is reduced by doing an EN regression, as proposed in Horvath’s paper, and then Bayesian learning is applied. This paper illustrates that this approach yields results that are on average superior to directly using BNN on the data, as well as using EN combined with BNN using other training algorithms, such as Levenberg-Marquard’s algorithm. The proposed approach is validated using real datasets that were obtained from GEO repository (https://www.ncbi.nlm.nih.gov/geo/). This method is implemented in an R function, which available through a package called `methylclock`.

## 2 Methods

### 2.1 Proposed approach

Herein, we describe the statistical methods that we used in the two steps performed to predict chronological age using methylation data. The method assumes that input data are CpGs probes containing beta values obtained from methylation DNA arrays (Illumina 27K or 450K).

#### 2.1.1 Elastic net

The elastic net is a robust algorithm, Hui and Hastie [17], for linear regression that has the interesting property of making some of the coefficients equal to zero reducing in practice the amount of independent variables in the model. If we assume that we have a dependent variable (*y*) and several independent variables (*X*), then an elastic net regression is done by finding the estimator that minimises this equation (Hui and Hastie [17]):

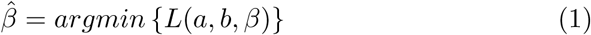

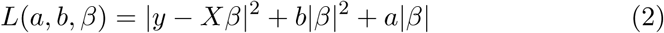

By doing this, the dimensionality of the problem is reduced. This helps us to deciding which independent variables to keep in the regression. This is particularly useful when there is a large number of independent variables and it is not clear which ones are relevant for the regression. However, keeping too many independent variables could cause the obtained expression to generalise poorly when new data is used.

#### 2.1.2 Levenberg Marquardt

Levenberg-Marquardt, Levenberg [18], Marquardt [19], “LM” is a commonly used training algorithm in neural networks, Hagan and Menhaj [20], Smaoui and Al-Yakoob [21], Bahram et al [22], Basterrech et al [23], that avoids calculating the Hessian matrix and has the goal of minimising a non-linear function. In mathematical terms, the goodness of the estimated *ŷ* values and the actual *y* values can be described using a chi distributed error that can be computed using the formula propose by Gavin [24]:

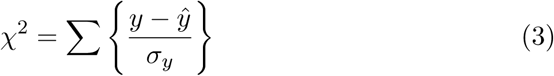

With the LM model commonly used as a training algorithm solving this equation (Gavin [24]):

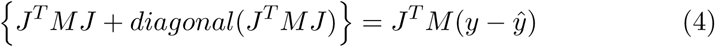

Where *ŷ* are the predicted values, *J* is the Jacobian and *M* is a diagonal matrix with each diagonal component equal to the inverse of the variance. The LM is a well-tested algorithm with application in fields as diverse as character recognition, as shown by Badi [25], or to model exponential increases of viruses, as shown by Novella [26]. For a more in depth description of the algorithm, we point the reader to the work by Chen and Gao [27].

#### 2.1.3 Bayesian regularisation

Another possible training algorithm for neural networks, which seems to have obtained better results when applied to the case of DNA methylation, is Bayesian regularisation. The purpose of Bayesian regularisation is, as described by Mackay [28], to “minimise a linear combination of squared errors and weights.” It is also designed to have good generalisation properties (it should be noted that this is the reference description given by some commercial software as Matlab). The issue of over-fitting is important while analyzing methylation in cells because the large amount of data makes it easy to fall due to the over-fitting issue. Over-fitting is, in simple words, an issue that arises where a neural network matches very closely the output of the training data but then produces poor results when applied to other data, which are not seen by the algorithm. In other words, the network does not generalise well.

It is common practice in this type of problem to try to minimise the sum of the squared errors:

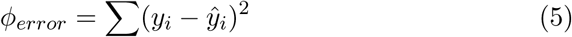

In Bayesian regularisation an additional term, Hagan and Foresee [29], the sum of the squares of the weights (*ϕ*_*weights*_) is added, with the expression becoming:

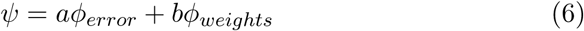

With the basic algorithm being (Hagan and Foresee, 1997):

1. Initialise parameters *a* and *b*
2. Calculate *ψ*
3. Using the approximation:

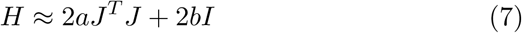 Estimate:

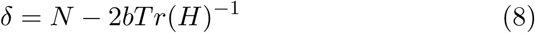
4. Estimate the value of the parameters:

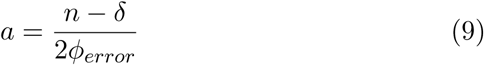

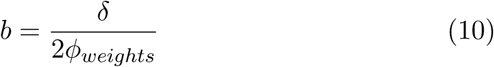
5. Repeat iteratively (starting after step 2)

For a more detailed explanation please see the original article (Hagan and Foresee [29]).

### 2.2 Trainning dataset

Horvath’s chronological age predictor was based on 8,000 samples from different tissues and cell types. Probes of these samples were generated from the Illumina 27K DNA methylation arrays. The age prediction was based on 353 CpGs that were selected using EN. Using these CPGs as the first step, we then trained a Bayesian Neural Network using data from 720 individuals, who were used to study the effect of ageing on methylation (GEO accession number GSE41037, https://www.ncbi.nlm.nih.gov/geo/). Methylation data was obtained from Illumina HumanMethylation27 bead chip that provides methylation levels across approximately 27,000 CpGs measured in blood. We used 15 percent of the data for internal testing purposes. Following standard procedures, these data were not used during the training phase to evaluate how the network generalised to new data and, more importantly, to assess the issue of over-fitting.

### 2.3 Validation datasets

We used 8 different GEO datasets measured in different tissues and platforms to assess the external validity of the proposed method as well as to compare model’s performance with Horvath’s method (Table 1).

**Table 1:** GEO datasets used for external validating purposes.

| **DNA origin** | **Platform** | **n** | **age range** | **samples** | **GEO number** |
| --- | --- | --- | --- | --- | --- |
| Whole Blood | 450K | 43 | (47, 59) | Healthy | GSE53128 |
| Whole Blood | 450K | 231 | (34, 72) | Cancer/Control | GSE49996 |
| Whole Blood | 450K | 214 | (51, 82) | Alcohol | GSE112987 |
| Whole Blood | 27k | 172 | (33, 80) | Healthy | GSE58045 |
| Blood | 27k | 214 | (42,93 | Healthy | GSE19711 |
| Blood | 450K | 16 | (21 32) | Healthy | GSE65638 |
| Blood PBMC | 27k | 91 | (24, 45) | Healthy | GSE37008 |
| Blood CD4+CD14 | 27k | 50 | (16, 69) | Healthy | GSE20242 |

### 2.4 Performance evaluation

We computed two statistics in the different GEO validation sets to evaluate the performance of our age predictors: (1) the correlation between predicted age and chronological age (R-squared); and (2) the Root Mean Square Error (RMSE). We then computed the standard error of the R-squared and RMSE using the formulas provided in [30] and [31], respectively. We meta-analyzed the difference of the R-squared and RMSE between our proposed method and Horvath’s approach using both fixed and random effect models implemented in `meta` R package [32].

## 3 Results

We first applied BNN to the data without reducing the dimensionality using EN (e.g. direct neural network approach). The network was trained 100 times using LM and another 100 times using Bayesian regularisation in both cases the results were less than optimal with substantial dispersion in out-of-sample values obtained for the R-squared of the regression comparing the predicted age estimates with the chronological age of the patients with the network in several cases, stopping after 1,000 iterations without reaching the target error. The input for the network in both cases was the methylation levels of the 27,000 CpGs, with no further transformation. The target output was the age of the patients. The network had two layers (only one hidden) and had 10 neurons in the hidden layer. The low R-squared values obtained from the simulations indicate that the model is not capturing too much variability (median: 0.34, range: 0.22, 0.83). These results suggests that applying a neural network directly to the data is not the best approach because the variance that is explained by this approach is too modest. This is likely to happen because of the issue of local minima in neural networks, which are of particular importance when, as in this the case, there are a large number of input data (CPGs) and a smaller number of samples (see Bo [33]). Reducing the dimensionality of the data can substantially increase the accuracy of the forecasts.

Our two steps approach aims to overcome this difficulty. We first reduce the dimensionality of the data by using EN and then apply a better predictor methodology (e.g. BNN). The EN step is performed by using the CpGs obtained in Horvath [1] that were obtained after regressing chronological and age on about 27,000 CpGs from 8,000 individuals. The CPGs selected in that paper have proven to be useful for age predicting purposes. Consequently, the dimensionality of the problem is reduced from 27,000 variables per individual to 353 variables. The methylation data for these CpGs were used as the input for BNN. The BNN chosen has two layers with 10 neurons in the hidden layer. The neural networks were then trained using both the Levenberg-Marquardt algorithm and Bayesian regularisation. The results of a neural network will change every time that it is trained because we used different initialisation parameters to avoid sub-optimal local convergence. We run 100 simulations for LM algorithm and another 100 for Bayesian regularisation. Figure 1 shows the distribution of R-squared values of each simulation for the two different training methods.

**Figure 1:**
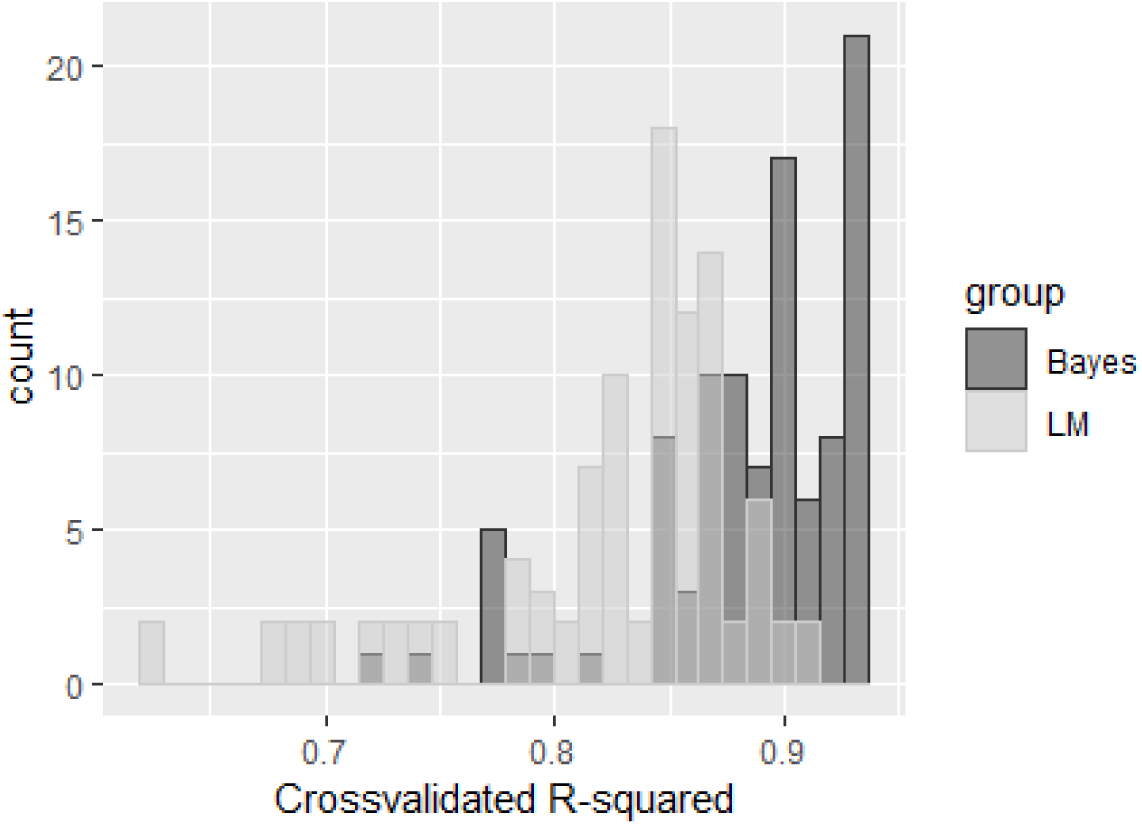
R-squared histograms for LM and Bayesian learning methods of simulated datasets.

We observed that Bayesian training clearly outperforms the LM method since the median R-squared for LM is 0.85 (range: 0.63 to 0.92) while for Bayesian training is 0.90 (range: 0.71 to 0.93) (Table 2). These observed differences are statistically significant after applying Wilcoxon test (*p* = 6.4 × 10^−53^). The same conclusion is obtained when comparing our two stage proposed method (EN + BNN) with one based on a BNN applied to the entire dataset (Wilcoxon test, *p* = 6.3 × 10^−28^). This indicates that reducing the dimensionality actually (statistically significantly) improved the forecasts.

**Table 2:** Median and range of R-squared values obtained from two Neural Networks trained using two methods different methods (Levenberg-Marquardt and Bayesian) after dimension reduction of one of the validation datasets (GSE58045) in 100 simulated datasets.

| Algorithm | R-squared |  |
| --- | --- | --- |
|  | Median | Range |
| LM | 0.8492 | [0.6274, 0.9137] |
| Bayesian | 0.900 | [0.7152, 0.9340] |

We applied our proposed method to the internal test dataset (15% out-of-sample of GEO number GSE41037 not using when creating the predictive model, n=108). The regression of the predicted values against the chronological age can be seen in Figure 2. We observe a good performance with an R-squared equal to 0.94, being 0.97 in the training data (85% of GSE41037 dataset, n=612), which obviously over-estimates the model’s predictive value.

**Figure 2:**
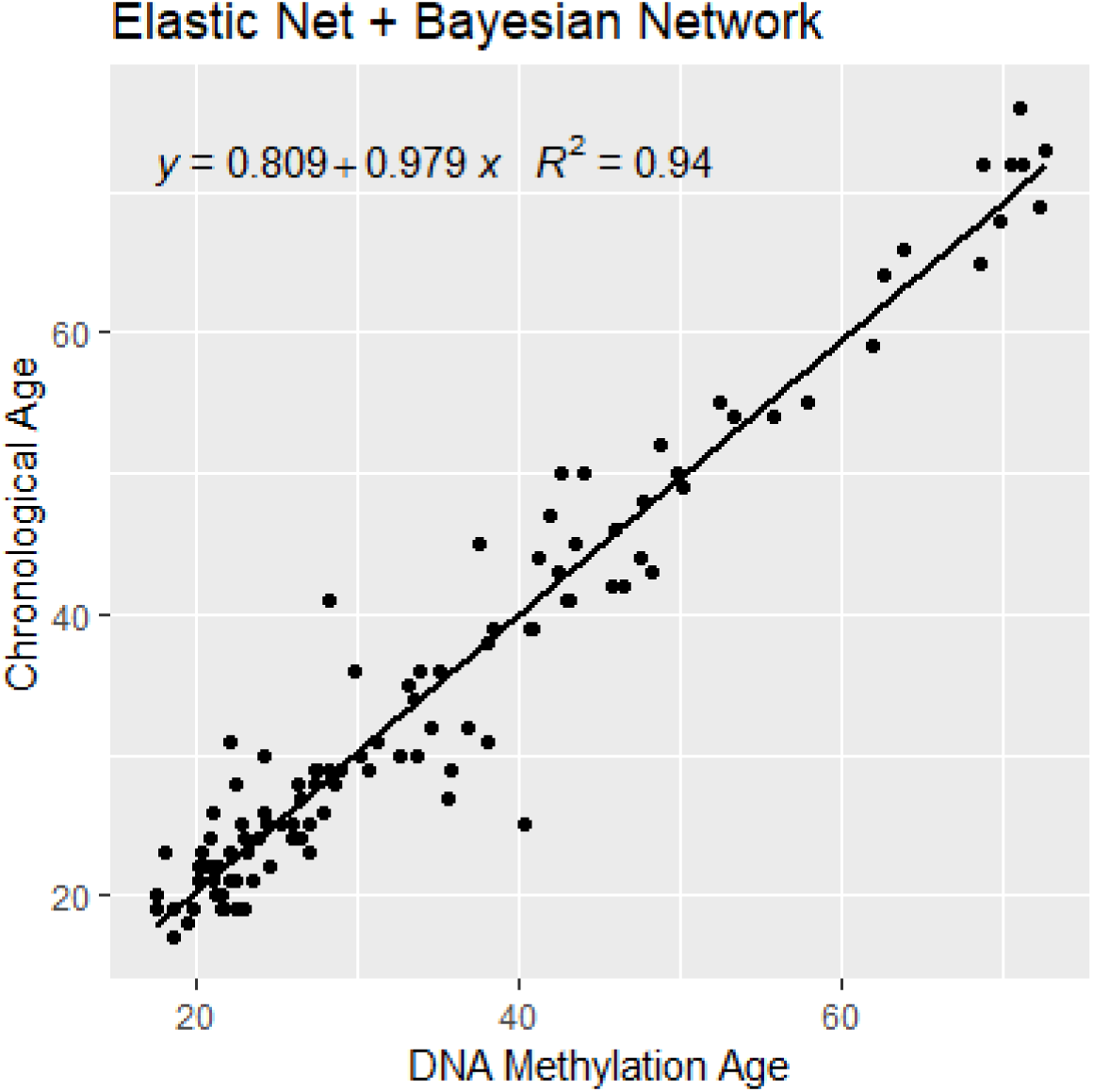
Regression of the predicted age against the chronological age using the internal test dataset (15% out-of-sample of the training dataset GSE41037).

### 3.1 Bayesian neural network implementation

The Bayesian neural network was trained using Matlab. The first step was to create a neural network structure of two layers, one of which is hidden. The hidden layer has 10 neurons. This structure generated better results than more complex neural networks with larger amount of neurons or of hidden layers. Some of the more complex networks analyzed generated good results for the databases analyzed but generalised poorly when handling new databases. The database GSE41037 was used for training purposes with 15 percent of the data kept aside for testing purposes. The training algorithm used was Bayesian regularisation, using the package `trainbr` in Matlab. A ten times cross validation approach was followed for training purposes. Consequently, a net structure was generated in Matlab. For convenience that net structure was transformed into a function using the Matlab function `genFunction`. This function takes as an input a matrix of values in which each column represents one patient and each row represents the methylation value for each CPG, as previously mentioned we used the CPGs obtained by Horvath [1]. The function then generates a vector output that is the obtained value for the age of each patient is then compared to the chronological age of the patient using simple linear regression. The function was then transformed into C++ code using the Matlab application “Matlab Coder”. An R function that calls this C++ code was created to allow users predicting age using DNA methylation data that is available at https://github.com/isglobal-brge/methylclock.

### 3.2 Validation

Our proposed method was tested using 8 independent GEO datasets. The results obtained from our two stage approach (Elastic Net + Bayesian Neural Network) were compared with those obtained using Horvath’s method. We selected some GEO datasets that have been published after Horvath’s paper was published to avoid including individuals that were used to build Horvath’s predictive model and, hence, avoid overfitting (Table 1). Figure 3 shows the meta-analysis of R-squared and RMSE obtained from our proposed method and Horvath’s approach. We observe that, as expected, there is a large heterogeity in the results (p of heterogeneity ¡0.01). There are some GEO datasets were both methods are providing similar R-squared (GSE20067, GSE51032 and GSE101764) but, in general, a better performance when using our proposed method is achieved (Figure 3A). In summary, we observe that our method explained a 4% more variability of the chronological age than Horvath’s method (CI95%: 2%-5%). Similar conclusion was achieved when comparing RMSE (Figure 3B). This is not surprising considering that there could be some degree of non-linearity in the relationship between methylation levels and aging. For processes that are, at least to some degree, non-linear, neural networks should provide better results than linear regression.

**Figure 3:**
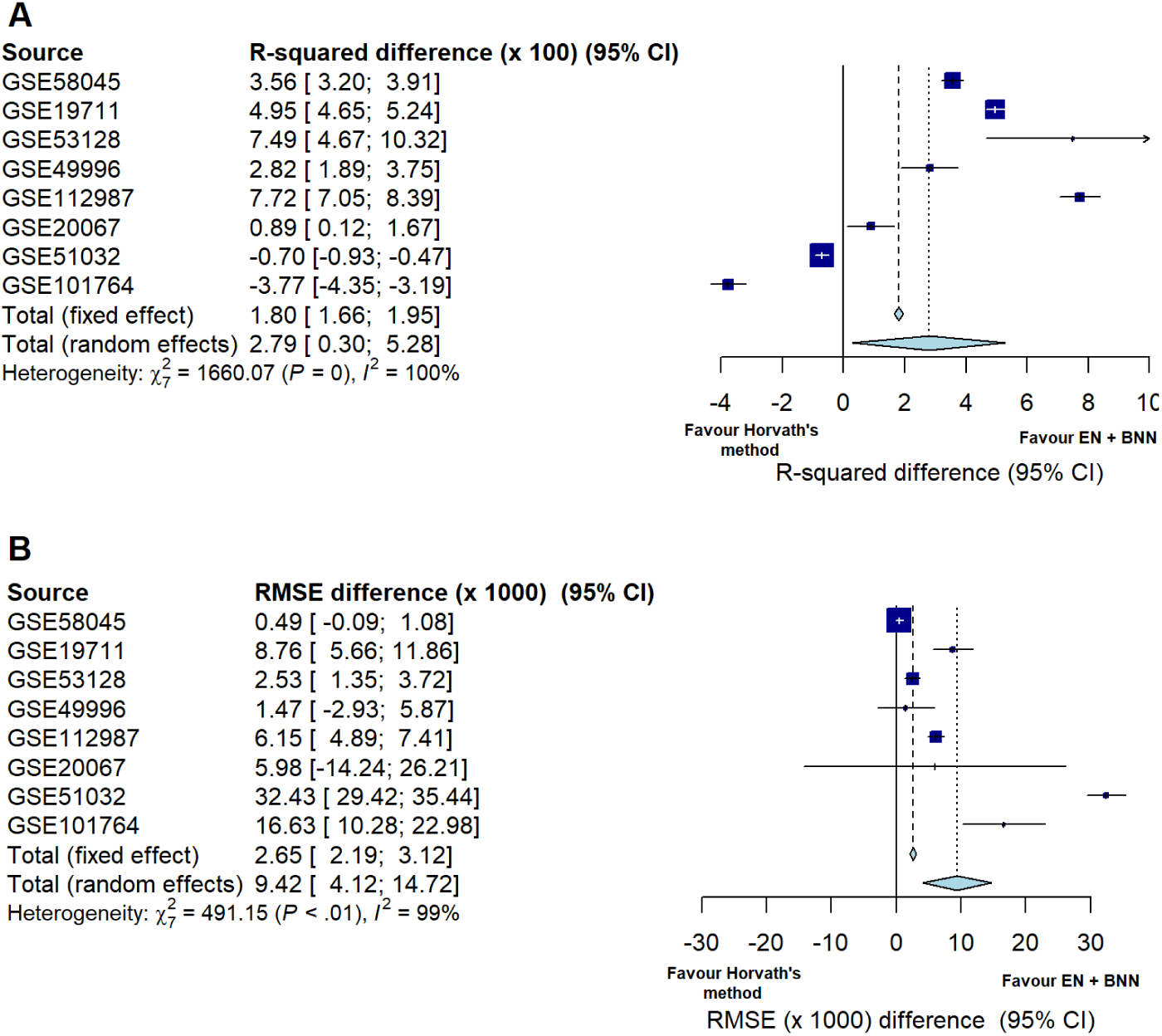
Validation results using different GEO datasets corresponding to our proposed method (Elastic Net + Bayesian Neural Networks – EN + BNN) and Horvath’s method. The panel A shows the difference of the R-squared obtained after regressing chronological age with the age predicted using our proposed method and Horvath’s approach. Panel B shows the same comparison for the RMSE (x1000).

## 4 Discussion

Biological clocks built using EN regression followed by neural networks trained with Bayesian regularisation applied to blood methylation data seem to produce better results than using a single step method (either using EN or BNN to the entire methylation data). Our conclusion is based, first, on the results obtained when comparing the estimated age with the chronological age of the patients using only EN or neural networks with other training algorithm, such as Levenberg-Marquardt’s algorithm. The second piece of evidence in favor of our proposed method relies on the comparisons performed with Horvath’s method in different real datasets.

One of the base assumptions is that there is some non-linearity in the relationship between methylation levels and aging that cannot be captured when using linear models, as in the case of Horvath’s method that uses elastic net regression models. Meanwhile, it is likely that the reason why reducing the dimensionality of the data before applying neural networks produces good results is related to the issue of local minima.

In conclusion, our results support the hypothesis that there is some degree on nonlinearity in the relationship of methylation levels and age. Some existing linear models predict age with a reasonable level of accuracy but (as shown in this article) neural networks, particularly after reducing the dimensionality of the data, generate better results. This suggests that there is some level on non-linearity in the process. Having an R implementation of our predictive model will help biomedical researchers to incorporate an epigenetic biomarker to assess its impact in age-related complex diseases, such as cancer or Alzheimer’s, among others.

## 5 Data accessibility

The biological clock can be accessed in: https://github.com/isglobal-brge/methylclock. All the data is accessible at GEO Database (https://www.ncbi.nlm.nih.gov/geo/) using the mentioned accessing codes.

## 6 Authors’ contributions

GA proposed the method to create the DNA methylation clock, created the C++ code of the Bayesian Network analyzed the data and wrote the manuscript. JRG validated the proposed model, created the R package, wrote the manuscript and oversaw the project. Both authors have read and approved the final version of the manuscript.

